# Generating spatiotemporal patterns of linearly polarised light at high frame rates for insect vision research

**DOI:** 10.1101/2022.01.31.478537

**Authors:** Jack A. Supple, Léandre Varennes-Phillit, Dexter Gajjar-Reid, Uroš Cerkvenik, Gregor Belušič, Holger G. Krapp

## Abstract

Polarisation vision is commonplace among invertebrates; however, most experiments focus on determining behavioural and/or neurophysiological responses to static polarised light sources rather than moving patterns of polarised light. To this end, we designed a polarisation stimulation device based on superimposing polarised and non-polarised images from two projectors, which can display moving patterns at frame rates exceeding invertebrate flicker fusion frequencies. A linear polariser fitted to one projector enables moving patterns of polarised light to be displayed, whilst the other projector contributes arbitrary intensities of non-polarised light to yield moving patterns with a defined polarisation and intensity contrast. To test the device, we measured receptive fields of polarisation sensitive *Argynnis paphia* butterfly photoreceptors for both non-polarised and polarised light. We then measured local motion sensitivities of the optic flow-sensitive lobula plate tangential cell H1 in *Calliphora vicina* blowflies under both polarised and non-polarised light, finding no polarisation sensitivity in this neuron.

## Introduction

Light becomes polarised when scattered by small particles or reflected from a surface (Cronin and Marshall, 2011). Polarisation refers to the distribution of electric field vector orientations within a beam of light. The angle and the degree (i.e., ratio of the polarised to total light intensity) of linear polarisation (AoP, and DoLP, respectively) are dependent on both the relative position of the light source and the physical structure of the polarising material (Labhart, 2016). Thus, alongside wavelength, intensity, and the propagation direction of light, AoP and DoLP provide information about the physical properties of the visual environment (Labhart, 2016). Consequently, many animals exploit patterns of linearly polarised light to guide tasks as diverse as navigation (Dacke, 2014; Zeil et al., 2014), communication (How et al., 2015; Mäthger et al., 2009; Sweeney et al., 2003), water-seeking (Obayashi et al., 2021), and object detection (Foster et al., 2014; Kelber et al., 2001; Meglič et al., 2019).

Historically, experimental polarised light stimuli usually comprised static light sources filtered through a linear polariser which is rotated throughout the experiment. This enabled investigation of widefield polarisation cues for navigation (Evangelista et al., 2014; Henze and Labhart, 2007; Mappes and Homberg, 2004; Mathejczyk and Wernet, 2019; Warren et al., 2018; Weir and Dickinson, 2012), polarisation sensitive photoreceptors (Bandai et al., 1992; Blum and Labhart, 2000; Weir et al., 2016; Zufall et al., 1989), and interneurons (Hardcastle et al., 2021; Heinze and Homberg, 2007; Träger and Homberg, 2011; Zittrell et al., 2020). However, this design is less suited for investigating the role of structured, spatiotemporal patterns of polarisation contrast in object detection and motion vision.

More recent solutions for generating moving patterns of polarisation contrasts involve modifying liquid crystal display (LCD) monitors, and have been used to investigate optomotor reflexes (Glantz and Schroeter, 2006) and object detection (How et al., 2014; Pignatelli et al., 2011; Temple et al., 2012). However, this LCD modification is restricted to displaying AoP contrasts within the range of rotation of the liquid crystals and is unable to generate polarised patterns over a non-polarised (and luminance equalized) background. An alternative method addressing these limitations utilized a digital light processing projector (DLP) to display video frames through a synchronously rotating linear polariser, relying on the low-pass temporal filtering properties of photoreceptors to integrate each pattern into a resultant polarisation contrast image (Stewart et al., 2017). However, the AoP resolution and frame rate of moving patterns are limited by the rotation speed of the polariser.

In many insects, the flicker fusion frequency, i.e. the frequency at which photoreceptor responses no longer discriminate between sequential contrast changes, falls between 100-300 Hz (Chatterjee et al., 2020; Cosens and Spatz, 1978; Ruck, 1961; Tatler et al., 2000). To generate stimuli that exceed these flicker fusion frequencies, we have designed a stimulation device based on superimposing polarised and non-polarised images from two high frame rate DLP projectors (Figure 1A). The superposition of images from the two DLPs enables the presentation of visual motion stimuli salient in either intensity-only contrast, polarisation-only contrast, or a combination of both intensity and polarisation contrasts. This design enables the DLPs to project motion images at high frames rates, delegating control of polarisation contrast to a dedicated projector.

**Figure 1:**
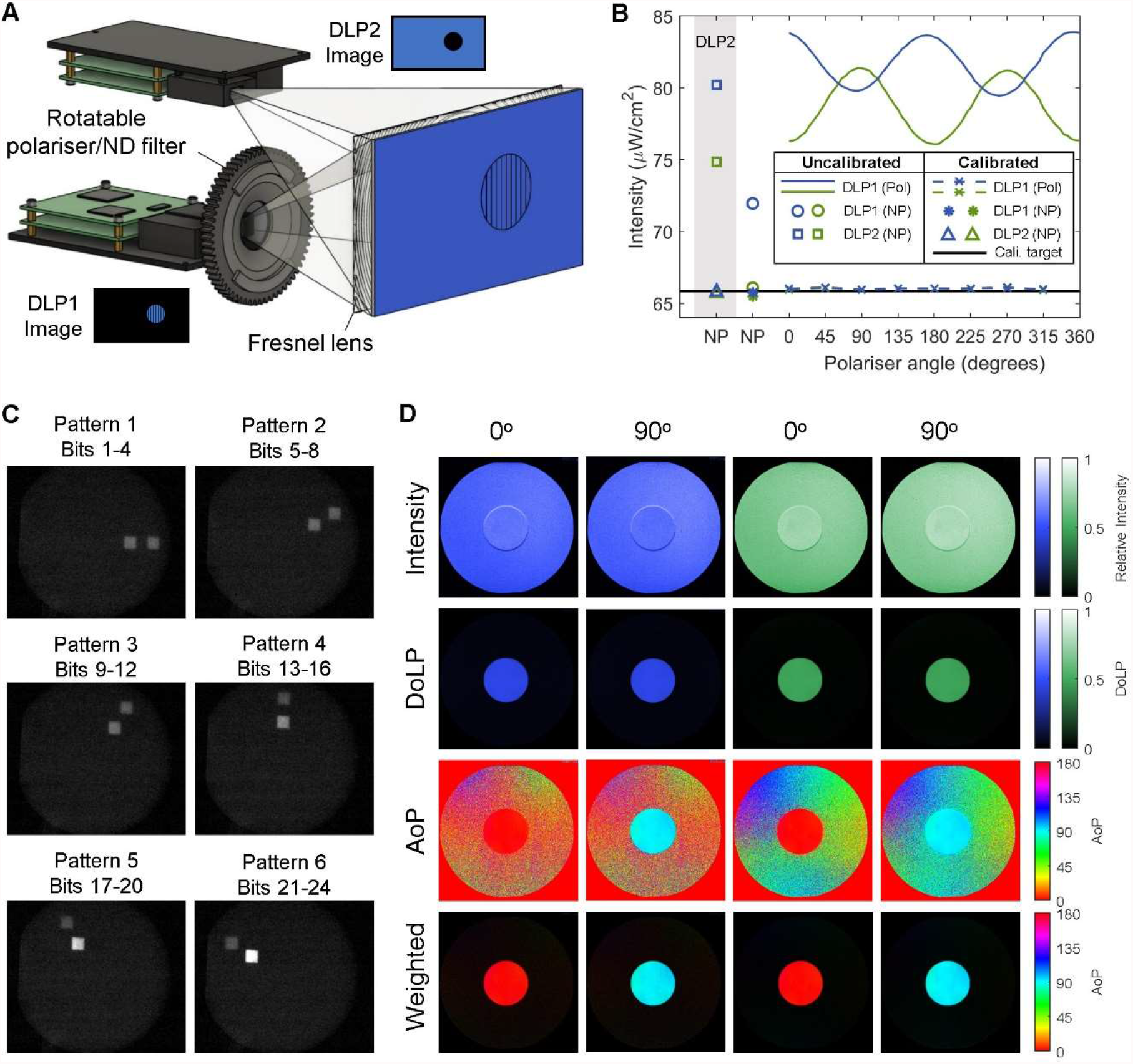
Dual DLP stimulation device for projecting moving patterns of polarised light. (A) Images from two DLP projectors are aligned and superimposed. Patterns can be projected with polarisation intensity contrasts (DLP1), or non-polarised intensity contrasts (DLP2). Superimposing inverted images from the two DLPs nullifies intensity contrasts. Polariser and neutral density (ND) filters can be exchanged for control experiments. (B) LED luminance equalization of the two projected images for blue and green light. Polarised DLP1 intensities varied sinusoidally with polariser angle due to intrinsic DLP polarisation. The lowest uncalibrated luminance (here, DLP1 green non-polarised) was designated the calibration target to which other intensities were matched by adjusting LED currents. (C) To verify temporal synchrony, two squares, one from each projector, were filmed moving along circular trajectories with offset radii. The six panels correspond to the motion of each square along its circular trajectory at 360 Hz. Angular correspondence of each square position indicates that the two projectors are temporally synchronized. (D) Polarimetry of the dual DLP system displaying a polarised dot at 0^°^ and 90^°^ AoP against a luminance-equalized background of the same color for blue and green light. Top row: Normalised total intensity. Second row: DoLP. Third row: AoP. Bottom row: AoP with pixel brightness weighted by the DoLP (low DoLP pixels appear darker than higher DoLP pixels).

To demonstrate the performance of the device, we measured receptive fields of *Argynnis paphia* butterfly photoreceptors for both non-polarised and polarised light. Butterfly blue and green photoreceptors are maximally sensitive to light polarised along the dorsal-ventral and medial-lateral axes, respectively, with a polarisation sensitivity ratio ∼2 (Bandai et al., 1992; Kelber et al., 2001). We measured the receptive fields of these polarisation sensitive photoreceptors to confirm the operation of the dual DLP polarisation device.

We then used the device to characterise the receptive field of the Lobula Plate Tangential Cell (LPTC) H1 in the blowfly *Calliphora vicina* for non-polarised and polarised light. LPTCs are a large class of interneurons in the lobula plate that integrate local motion information across large regions of visual space, resulting in receptive fields selective for specific patterns of optic flow (Borst et al., 2010; Franz and Krapp, 2000; Hausen, 1984). H1 is a spiking LPTC sensitive to horizontal back-to-front optic flow across the ipsilateral eye equator, and projects to LPTCs in the contralateral lobula plate (Hausen, 1976; Horstmann et al., 2000; Krapp et al., 2001; Weber et al., 2010). The multiplication of time-delayed responses from neighbouring ommatidia renders the insect elementary motion detector dependent on the square of the intensity contrast (Buchner, 1976, 1984). Consequently, LPTCs are highly sensitive to changes in image contrast, making them well suited to test for intensity contrast artefacts in the superimposed, intensity-masked DLP images.

A subset of LPTCs synapse with descending neurons that control optomotor stabilisation reflexes in response to optic flow (Haag et al., 2007; Suver et al., 2016; Wertz et al., 2009a, 2009b). The optomotor response is driven primarily by broadband spectral sensitivity of R1-R6 photoreceptors (Yamaguchi et al., 2008). However, R7-R8 photoreceptors have been found to improve optomotor related motion detection via electrical coupling with R1-R6 photoreceptors in *Drosophila* (Shaw et al., 1989; Wardill et al., 2012). In *Tabanus* horseflies, a subset of R7-R8 photoreceptors within ‘pale’ type ommatidia comprise a blue-UV polarisation analyser (Meglič et al., 2019). Thus, if this R7-R8 polarisation analyser is present among other dipterans, we may expect to find polarisation sensitivity in LPTCs. However, in this study we test the H1-cell for polarisation sensitivity in *Calliphora vicina*, but do not find evidence for polarisation sensitivity.

## Methods and Materials

### Dual-DLP stimulation device

Two LightCrafter 3000 DLPs (Texas Instruments, USA) were mounted on a 2020 aluminium extrusion frame, projecting through a polarisation-preserving rear-projection screen (ST-Pro-X, Screen-Tech e.K., Hohenaspe, Germany). A Fresnel lens (Moxs, UK) was positioned behind the projection screen to collimate light from each DLP, reducing intensity gradients across each projected image. The top DLP (DLP2) was rotated upside-down and vertically displaced from the lower DLP (DLP1) until the projected images were superimposed (Figure 1A). To precisely align the two projectors, the top DLP was attached to the frame via a custom four-screw spring platform enabling adjustments in roll, pitch, and up-down translation (Figure S1A-B). Sideways and back-to-front translation and yaw rotation was enabled by a circular attachment of the spring platform to the extrusion beam (Figure S1C).

The bottom DLP1 was fitted with a 3D printed rotation mount (https://www.thingiverse.com/thing:5224352) controlled by an Arduino Nano, A4988 driver, and stepper motor (Nema 17HS4023). Small cylindrical neodymium magnets (6mm diameter x 2mm, first4magnets, UK) on the rotation mount enabled exchange of either a linear polariser (LPVISE2X2, Thorlabs, UK) or a non-polarising neutral density filter (Thorlabs, product NE05B). The top DLP2 was fitted with a fixed non-polarising neutral density filter (Thorlabs, product NE05B) to roughly equalise the luminance of the two DLPs. DLP intensities were precisely equalised by adjusting the current supply to each DLP LED (Figure 1B). Projection intensities were measured with a photodiode (RS Stock #: 642-4430) calibrated by a photometer (AccuPRO XP-4000 Plus, Spectronics Corporation, USA). Due to intrinsic polarisation of the system, LED luminance varied sinusoidally with rotation of the polariser (Figure 1B). This was compensated for by selecting the lowest uncalibrated luminance for all LEDs under both non-polarised and polarised conditions as the calibration target. An automated gradient descent algorithm adjusted each LED current to reach this luminance target, which were then stored as a polariser angle-dependent look-up table for each LED.

The two DLPs were connected to the display ports of an AMD RX 580 graphics card controlled by a ROG STRIX x470-f gaming motherboard, Ryzen 5 3600x CPU, and Corsair Vengeance LPX 2×8GB DDR4 3200 MHz RAM. DLPs were configured as an extended ‘Eyefinity display’ video wall with V-Sync enabled using the AMD Adrenaline graphics driver. Images were streamed to each DLP as 60Hz 24-bit RGB. GIF videos using StimulateOpenGL (StimGL) II v.20160216 (https://github.com/cculianu/StimulateOpenGL_II). Each DLP was configured to parse the 24-bit RGB images in BRG order as 4-bit sequences of information per pixel to yield a moving pattern frame rate of 360Hz. Temporal synchrony of the two projectors was verified by highspeed videography (1000 fps; SA3 Fastcam, Photron, UK), filming two test targets, one from each projector, moving along synchronised circular trajectories with offset radii (Figure 1C). Visual stimuli were synchronised with electrophysiology traces via a photodiode sensing a small 50×50 pixel box alternating black and white every frame.

### Polarimetric Imaging

Polarimetry of the dual DLP setup demonstrated the confinement of linear polarisation to the polarised DLP (Figure 1D). Images were calculated from a series of digital photographs taken through a camera-mounted circular polariser filter (i.e., distinct from the linear polariser mounted on the polarised DLP) with the angle of polarisation rotated in 45° increments using an automated polarimeter attachment (https://www.thingiverse.com/thing:5220854). Photographs were acquired on a tripod-mounted Canon 7D DSLR in manual mode (aperture f/8, shutter speed 1/10 seconds, ISO 100) using a 50 mm prime lens (Canon EF 50 mm f/1.8 STM) with an attached circular polariser filter (Hoya Pro1 Digital Filter, Japan). Images were acquired as. CR2 raw image files, then reformatted as 16-bit, un-brightened, and un-gamma-corrected TIFF files using DCRAW software (https://www.dechifro.org/dcraw/) to preserve linearity between camera sensor readings and light intensity (Sumner, 2013). Bayer mosaic RGGB color channels were kept separate, with the blue and a single green Bayer channel selected for analysis of the blue and green LEDs, respectively. AoP and DoLP were calculated from Stokes parameters S_0_-S_2_ as described previously (Foster et al., 2014, 2018):

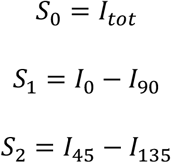

Where I_tot_ is the total intensity summed across images per pixel, and I_0_-I_135_ are the per pixel intensity measurements with the camera polariser positioned at 0° through 135° respectively. The angle of polarization, AoP, was calculated as:

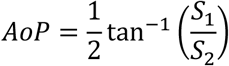

The degree of linear polarization, DoLP, was calculated as:

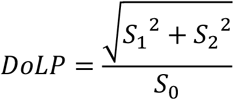

### Butterfly photoreceptor electrophysiology and receptive field mapping

*Argynnis paphia* butterflies were wild caught in Ljubljana, Slovenia in August 2021. Six photoreceptors from three *Argynnis paphia* adult males were recorded intracellularly using sharp borosilicate microelectrodes, filled with 3M KCl (R=80 MΩ). Recording electrodes were advanced through a small hole cut in the dorsal aspect of the eye; a 50 μm Ag/AgCl wire in the opposite eye served as a reference. Details of preparation and positioning are described elsewhere (Belušič et al., 2021). Signals were amplified in bridge mode through a SEC-10LX amplifier (NPI, Germany) and digitised at 30 kHz on a USB-6215 DAQ (National Instruments, USA). Data was collected and analysed in MATLAB.

First, photoreceptor spectral sensitivity was determined using flash stimulation from a LED spectral array, combined with a diffraction grating (Belušič et al., 2016, 2021). Photoreceptor receptive fields were then mapped by measuring membrane depolarisations in response to a series of 400 small (0.5°x0.5°) stationary square objects presented for 100ms at adjacent locations on a 20×20 grid, with 50ms delay between each square. Stimuli comprised of bright non-polarised squares on a dark background (Weber contrast: +1), bright polarised squares on a dark background (Weber contrast: +1), and polarised squares on a bright non-polarised background (Weber contrast: 0; Figure 2). 2D photoreceptor receptive fields for each condition were calculated from the mean photoreceptor voltage during object presentation relative to the baseline membrane potential for each stimulated 2D location (Figure S2A-B). Receptive fields were interpolated with a 2D cubic spline, smoothened with a 1 standard deviation 2D gaussian kernel, and normalised to the maximum response for each cell across stimulus trials (Figure S2C-D). For each cell, receptive field xy displacements were calculated from the non-polarised condition as the centre-of-mass of binary receptive fields thresholded at 30% of the maximum value. All receptive fields were offset by this non-polarised centroid coordinate prior to averaging. Polarisation tuning curves were calculated from the maximum receptive field responses (Figure S2C-D). Polarisation sensitivity ratios were calculated as the ratio between the maximum and minimum of the polarisation tuning curve.

**Figure 2:**
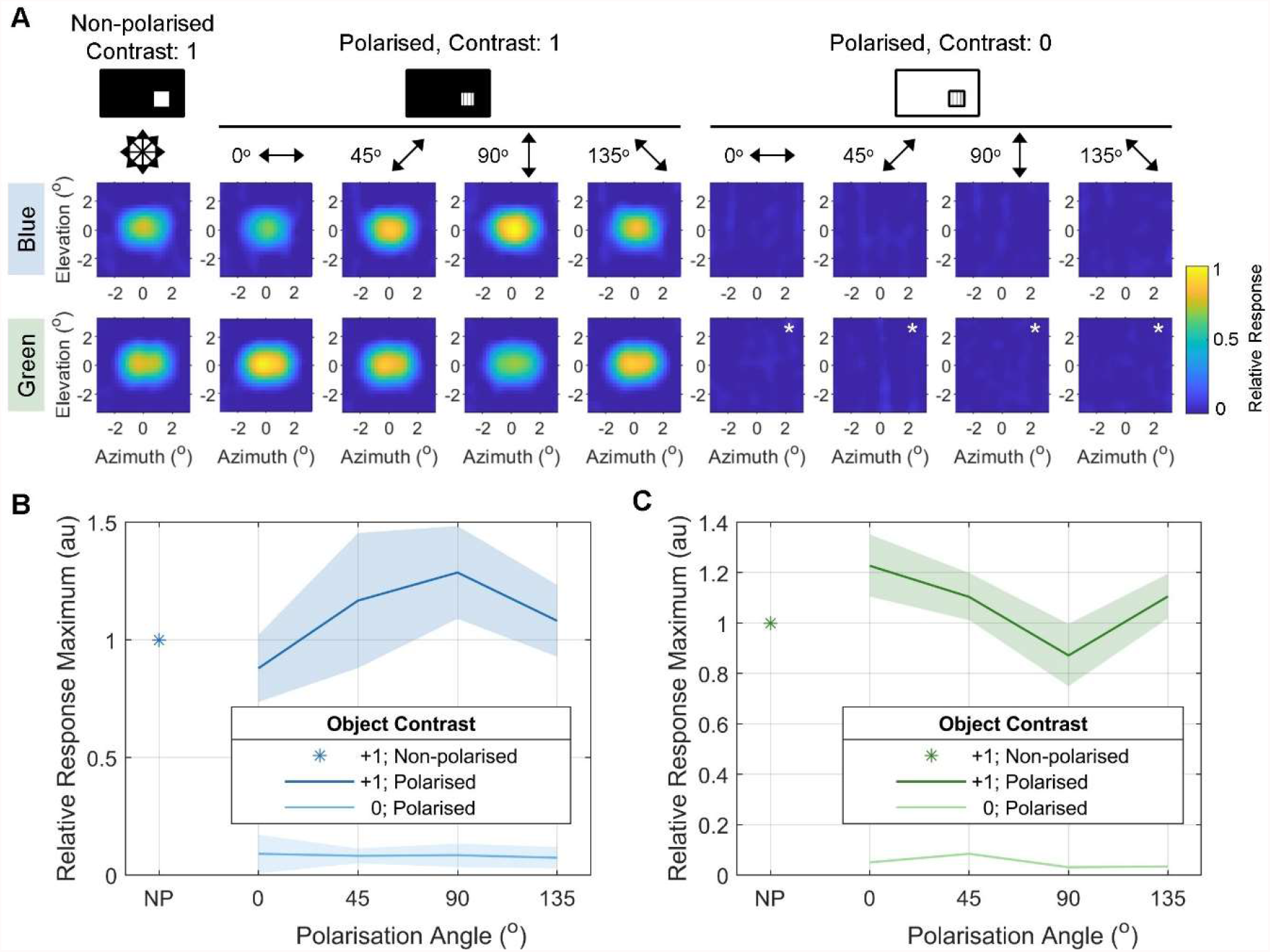
*Argynnis paphia* photoreceptor receptive fields for polarised and non-polarised light. (A) Receptive fields of polarisation-sensitive blue and green photoreceptors. Receptive fields were measured using non-polarised and polarised bright squares on a dark background (contrast=+1), and polarised objects masked with a bright non-polarised background (contrast=0). Blue: *N*=3 cells from 2 animals. Green: *N*=3 cells from 2 animals, except for masked polarised objects (*N*=1; asterisks). (B) Receptive field response maxima (see Figure S2) for blue-sensitive photoreceptors, normalized to non-polarised (NP) receptive fields (asterisk). Symbols/lines represent the mean of single trials per cell across animals, shaded regions represent 1 std. (C) Receptive field response maxima for green-sensitive photoreceptors, as in (B). Note *N*=1 for masked polarised objects (contrast: 0), so there is no standard deviation.

### Lobula Plate Tangential Cell (LPTC) electrophysiology and receptive field mapping

*Calliphora vicina* were acquired from a laboratory colony reared at 23°C and 60% humidity on a diet of water, sugar cubes, and pork liver. Nine *Calliphora vicina* adult males were mounted on a motorised mini– Goniometric Recording Platform (mini-GRP) (Huang et al. in preparation) 23cm from the dual DLP projection display. A sharp 3 MOhm tungsten electrode (UEWSHGSE3P1M, FHC, USA) was inserted into the exposed right-hand side lobula plate from the posterior aspect of the head capsule. Extracellular electrophysiological signals were amplified through a non-inverting operational amplifier at x10k gain (Huang and Krapp, 2013), and digitised at 30kHz on a NI USB-6215 DAQ (National Instruments, USA). Data was collected in MATLAB (Mathworks, USA). Extracellular action potential (spike) times were detected offline in Spike2 software (CED, UK) using a manually adjusted voltage threshold. Clusters of spike units were isolated manually after projecting the spike waveforms onto their first three principal components. In all cases, a single unit was detectable in the raw trace.

LPTC local motion sensitivities (LMS) and local preferred directions (LPD) were calculated using a fast stimulation protocol (Krapp and Hengstenberg, 1997) (Figure S3). A 7.6° circular dot rotated along a circular path of 10.4° diameter at 2 cycles per second for 10 cycles, repeated clockwise and counter-clockwise to cancel phase shifts introduced by the neuronal response latency. LPD was calculated as tangent to the circular mean of spike-triggered angular positions for clockwise (LPD_CW_) and counter-clockwise (LPD_CCW_) dots separately (Figure S3D). Due to the delay in neuronal response, spikes occur at a slight later phase in the stimulus after the preferred direction of motion (Krapp and Hengstenberg, 1997). The latency corrected LPD was calculated as the resultant vector of LPD_CW_ and LPD_CCW_ (Figure S3D). Local motion sensitivity (LMS) was calculated from clockwise and counter-clockwise dot responses as the mean difference in spike rate ±45° from the LPD (a_CW_ and a_CCW_; Figure 3D) and ±45° from the anti-LPD (b_CW_ and b_CCW_); i.e. (see Figure S3D):

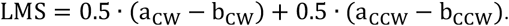

**Figure 3:**
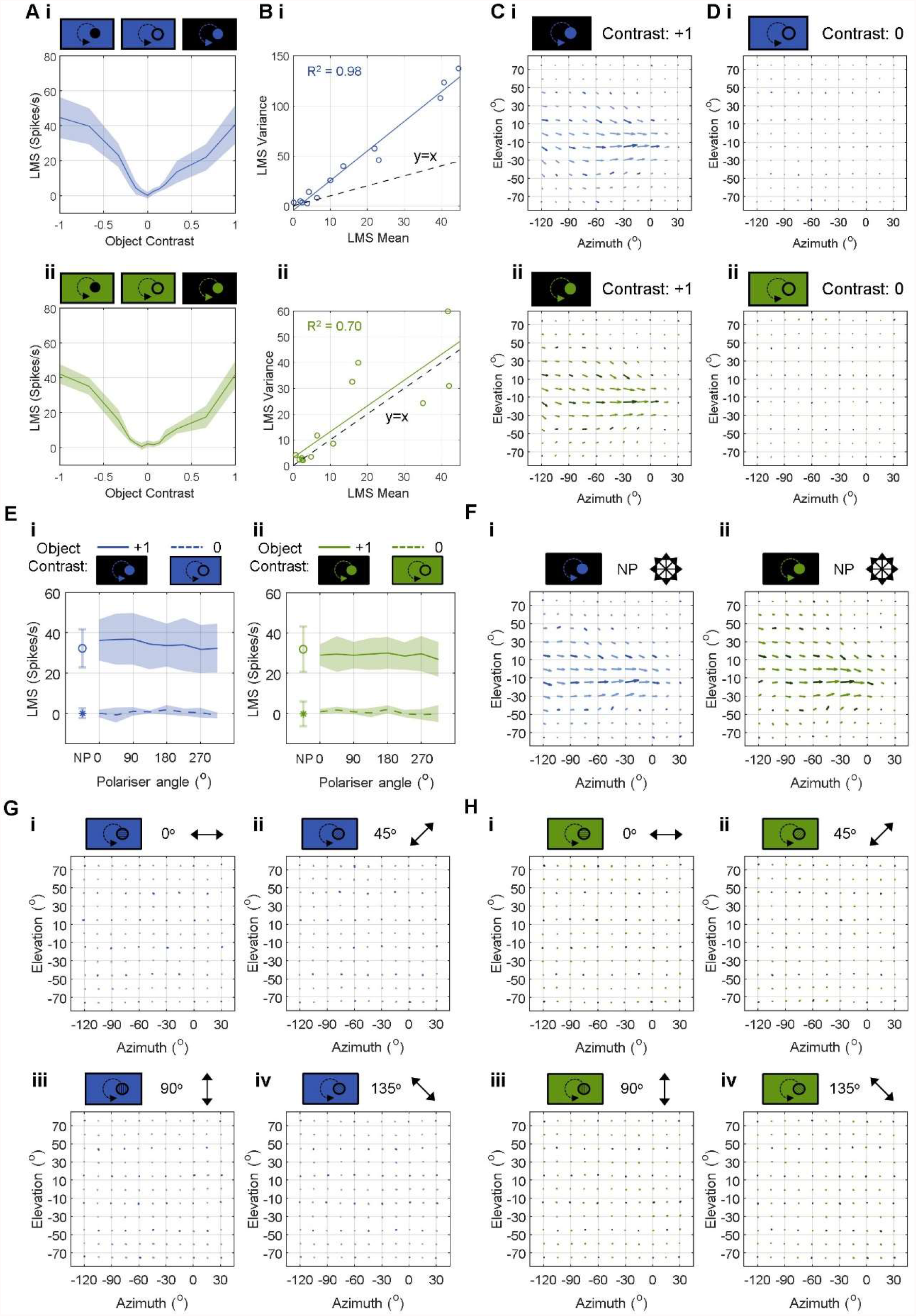
*Calliphora vicina* H1-cell motion responses to contrast artefacts and polarised light (*continued*) Figure 3 (*continued*) (A) H1-cell non-polarised control contrast tuning curves for (i) blue and (ii) green light. Both projectors are non-polarised. Contrast is varied from dark object/bright background (Weber contrast=-1), through to bright object/bright background (Weber contrast=0, i.e., intensity-nullified control condition), through to bright object/dark background (Weber contrast=+1). Data plotted as mean ± 1 std. *N*=7 animals. (B) H1-cell non-polarised LMS spike rate variance plotted against mean. Same data as (E). Black dashed line represents y=x. (C) Average H1-cell receptive fields for non-polarised bright (contrast=+1) (i) blue and (ii) green moving dots. Positive elevation values correspond to the dorsal visual field. Positive azimuth values correspond to the right visual field. Vector direction represents local preferred direction (LPD), vector length represents relative local motion sensitivity (LMS). Dark vectors represent sampled positions; lighter vectors are interpolated. Blue: *N*=6 animals; Green: *N*=5 animals. (D) Average H1-cell receptive fields for non-polarised and intensity contrast-nullified (contrast=0) (i) blue and (ii) green moving dots. Format same as (F). Blue: *N*=6 animals; Green: *N*=5 animals. (E) H1-cell polarisation tuning curves for (i) blue and (ii) green light. Bright (contrast=+1) and intensity contrast-nullified (contrast=0) dots were presented for non-polarised (NP) and polarised conditions at 45° AoP increments. 0° AoP aligns with the eye equator. Circle and solid line represent contrast=+1. Asterisk and dashed line represent contrast=0. Data plotted as mean ± 1 std. (F) Average H1-cell receptive fields of cells in (G-H) when stimulated with non-polarised bright dots (contrast=+1) for (i) blue and (ii) green light. Vector plot as in (C). *N*=4 animals for each spectral condition. (G-H) Average H1-cell receptive fields of cells in (F) when stimulated with polarised and intensity contrast-nullified (contrast=0) dots at 45° AoP increments for (G) blue and (H) green light. Vector plot as in (C). *N*=4 animals for each spectral condition.

Non-polarised contrast tuning curves (Figure 3A) were measured by varying the greyscale intensities of the moving dot and background, from a dark dot on a bright background (Weber contrast=-1), through to a bright dot on a bright background (Weber contrast=0), to a bright dot on a dark background (Weber contrast=+1). Polarisation tuning curves (Figure 3E) were measured for contrasts of 0 (bright dot on a bright background) and +1 (bright dot on a dark background) for 45° AoP increments through two full rotations of the polariser. LMS values were first averaged over trials within individual animals, then averaged across animals. Both non-polarised control contrast tuning curves and polarisation tuning curves were measured at a single location corresponding to the receptive field location with maximum LMS.

All LPTC receptive fields (Figure 3C-D, F-H) were characterised by measuring the LMS and LPD across a series of positions within the left and frontal visual field extending from -120° to +30° azimuth, and -75° to +75° elevation. Flies were automatically rotated between each stimulus presentation using the motorised mini-GRP. Vector fields were interpolated up to 15° increments across the sampled spatial range using a cubic spline. Receptive field vectors were normalised to the maximum value in the non-polarised condition prior to averaging across animals.

## Results and Discussion

### *Argynnis paphia* photoreceptor receptive field characterisation with polarised light

To test whether the device elicits polarisation sensitive neuronal responses, we recorded intracellularly from polarisation-sensitive photoreceptors in the retina of three *Argynnis paphia* adult male butterflies (Figure 2). Polarisation-sensitive *Argynnis paphia* photoreceptor receptive fields resembled 2D-gaussians for both non-polarised and polarised objects on dark backgrounds (Figure 2A). Peak receptive field responses were maximal for AoP of 90° (i.e., dorsal-ventral axis) and 0° (aligned with the eye equator) for blue and green-sensitive photoreceptors, respectively (Figure 2B-C). Polarisation sensitivity (PS) ratios were 1.5±0.2 (*N*=3) for blue photoreceptors, and 1.4±0.4 (*N*=3) for green photoreceptors. These PS values were lower than the ratio of ∼2 reported previously (Bandai et al., 1992; Kelber et al., 2001), which may have resulted due to a lower DoLP of light after scattering through the projection screen compared to un-scattered light used in previous experiments. Response magnitudes for non-polarised objects were intermediate between the maximal/minimal polarisation responses (Figure 2B-C), consistent with intermediate rhabdom photon capture for non-polarised light compared with polarised light of equal luminance. Overall, the dual-DLP polarisation display is sufficient to elicit polarisation-dependent responses in butterfly photoreceptors.

Photoreceptors did not respond to polarised objects upon a non-polarised intensity-masked background (Figure 2A-C), implying that the photon capture contrast between a polarised object and non-polarised background of equal luminance is insufficient to stimulate changes in photoreceptor potential. This suggests that moving patterns of polarisation-only contrast remain invisible, and some combination of polarisation and intensity contrasts are required for polarisation vision. Indeed, this agrees with the proposed integration of polarisation and intensity cues determined from the horsefly retina (Meglič et al., 2019). However, it remains possible that small photoreceptor depolarisations could be spatiotemporally pooled, with downstream neurons averaging and correlating responses across the retina.

### *Calliphora vicina* H1-cell is insensitive to superimposed projection artefacts

Despite best efforts to align the two superimposed images and equalise screen luminance, contrast artefacts between the two images remain detectable by the human eye (Figure 1D). To test whether these contrast artefacts are detectable by insects, we recorded extracellular responses from *Calliphora vicina* H1-cell under a non-polarised control condition in which both DLPs were fitted with neutral density filters (Figure 3A-D). Superimposed image contrasts were varied from a dark object/bright background (contrast=-1), through to a bright object/bright background (contrast=0), through to a bright object/dark background (contrast=+1). At 0 contrast, the two DLP images should cancel out, resulting in a featureless projection. Neuronal responses evoked under these conditions therefore arise from intensity contrast artefacts (e.g., mismatched luminance or positional misalignment between DLPs).

H1-cell LMS contrast tuning curves were V-shaped, reaching a minimum LMS of 0.2±1.9 spikes/s at 0 contrast for blue light (*N*=7; Figure 3Ai), and 0.7±2.0 spikes/s at -0.07 contrast for green light (*N*=7; Figure 3Aii). LMS variance increased linearly with mean LMS, albeit at a higher rate for blue compared with green light (Figure 3B). H1-cell receptive field structures agreed with the LPD and LMS distributions previously obtained using electromechanical stimuli (Krapp and Hengstenberg, 1997; Krapp et al., 2001). There were no differences between H1-cell receptive fields when stimulated with non-polarised blue or green light (Figure 3C). There were no responses across the H1-cell receptive field region for the non-polarised intensity contrast-nullified condition for both blue and green light (Figure 3D), suggesting that contrast artefacts were undetectable across the H1-cell receptive field.

The steeper rate of LMS variance increase with LMS mean for blue light (Figure 3B) suggests that motion vision is noisier when limited to this spectrum. This may reflect input from noisy blue-sensitive photoreceptor channels, which is only detectable when stimulated with blue light. In Diptera, pale ommatidia R8 (R8p) photoreceptors express blue-sensitive Rh5 opsin, and have noisier receptor potentials compared with R1-6 photoreceptors (Meglič et al., 2019). In *Drosophila*, R7-R8 photoreceptors form electrical synapses with R1-R6 photoreceptors, contributing to motion vision (Wardill et al., 2012). R8p are therefore candidates for this additional noisy input to the H1-cell and supports the hypothesis that multiple spectral inputs converge on dipteran motion vision pathways.

### *Calliphora vicina* H1-cell is insensitive to the angle of polarised light

To test *Calliphora vicina* H1-cell polarisation sensitivity, we presented bright (contrast=+1) and intensity contrast-nullified (contrast=0) moving dots at 45° AoP increments in the centre of the receptive field. LMS responses were not modulated by AoP in either contrast condition (Figure 4E). Similarly, there were no responses to intensity contrast-nullified polarised dots when presented throughout the H1-cell receptive field (Figure 4G-H), suggesting *Calliphora vicina* H1-cell is insensitive to polarised light.

The polarisation insensitivity of *Calliphora vicina* H1-cell implies that (i) *Calliphora* photoreceptors are insensitive to polarised blue or green light; and/or (ii) polarisation sensitive photoreceptors do not contribute to H1-cell motion sensitivity. In *Tabanus*, R8p photoreceptors are responsible for blue light polarisation sensitivity (Meglič et al., 2019). The contribution of R8-R7 to motion vision in *Drosophila* (Wardill et al., 2012), in addition to our putative finding that R8p contributes to H1-cell motion sensitivity (Figure 3B), suggests that H1-cell receives input from multiple photoreceptor types. Therefore, H1-cell polarisation insensitivity likely reflects photoreceptor polarisation insensitivity. Reproducing these polarisation motion sensitivity experiments in *Tabanus*, in which photoreceptor blue polarisation sensitivity is established (Meglič et al., 2019), will allow us to test this hypothesis.

## Conclusion

We have presented and tested a versatile device for projecting moving patterns of polarised light at high frame rates for insect vision, using two superimposed DLP projections. The device stimulates AoP-dependent responses in butterfly photoreceptors, and experiments in blowfly H1-cell suggest that residual contrast artefacts in the superimposed projection do not elicit motion sensitive responses. Furthermore, no evidence was found for polarisation sensitivity in blowfly H1-cell.

## Author Contributions

HGK and GB acquired financial support for the project. HGK conceived of the project. JAS, LVP, DGR built the polarisation device. HGK, JAS, LVP, UC, and GB designed the experiments. JAS and GB collected the data. JAS analysed the data and wrote the first draft; all authors contributed to the final version.

## Acknowledgments

We are grateful to the Air Force Research Laboratory for financially supporting this work. Thanks to all members of the Department of Biology, University of Ljubljana for their hospitality and offer of laboratory space over summer 2021. Thanks to Jiaqi Huang, Doekele Stavenga, Mathias Wernet, Nick Roberts, Primož Pirih, and Ric Wehling for their insight and comments throughout the project.

## References

Bandai, K., Arikawa, K., and Eguchi, E. (1992). Localization of spectral receptors in the ommatidium of butterfly compound eye determined by polarization sensitivity. J. Comp. Physiol. A 171, 289–297.

Belušič, G., Ilić, M., Meglič, A., and Pirih, P. (2016). A fast multispectral light synthesiser based on LEDs and a diffraction grating. Sci. Rep. 6, 1–9.

Belušič, G., Ilić, M., Meglič, A., and Pirih, P. (2021). Red-green opponency in the long visual fibre photoreceptors of brushfoot butterflies (Nymphalidae). Proc. R. Soc. B 288.

Blum, M., and Labhart, T. (2000). Photoreceptor visual fields, ommatidial array, and receptor axon projections in the polarisation-sensitive dorsal rim area of the cricket compound eye. J. Comp. Physiol. - A Sensory, Neural, Behav. Physiol. 186, 119–128.

Borst, A., Haag, J., and Reiff, D.F. (2010). Fly Motion Vision. Annu. Rev. Neurosci. 33, 49–70.

Buchner, E. (1976). Elementary movement detectors in an insect visual system. Biol. Cybern. 24, 85–101.

Buchner, E. (1984). Behavioural Analysis of Spatial Vision in Insects. In Photoreception and Vision in Invertebrates, (Springer US), pp. 561–621.

Chatterjee, P., Mohan, U., Krishnan, A., and Sane, S.P. (2020). Evolutionary constraints on flicker fusion frequency in Lepidoptera. J. Comp. Physiol. A Neuroethol. Sensory, Neural, Behav. Physiol. 206, 671–681.

Cosens, D., and Spatz, H.C. (1978). Flicker fusion studies in the lamina and receptor region of the Drosophila eye. J. Insect Physiol. 24, 587–594.

Cronin, T.W., and Marshall, J. (2011). Patterns and properties of polarized light in air and water. Philos. Trans. R. Soc. B Biol. Sci. 366, 619–626.

Dacke, M. (2014). Polarized light orientation in ball-rolling dung beetles. In Polarized Light and Polarization Vision in Animal Sciences, Second Edition, (Springer Berlin Heidelberg), pp. 27–39.

Evangelista, C., Kraft, P., Dacke, M., Labhart, T., and Srinivasan, M. V. (2014). Honeybee navigation: critically examining the role of the polarization compass. Philos. Trans. R. Soc. B Biol. Sci. 369.

Foster, J.J., Sharkey, C.R., Gaworska, A.V.A., Roberts, N.W., Whitney, H.M., and Partridge, J.C. (2014). Bumblebees learn polarization patterns. Curr. Biol. 24, 1415–1420.

Foster, J.J., Temple, S.E., How, M.J., Daly, I.M., Sharkey, C.R., Wilby, D., and Roberts, N.W. (2018). Polarisation vision: overcoming challenges of working with a property of light we barely see. Sci. Nat. 105, 1–26.

Franz, M.O., and Krapp, H.G. (2000). Wide-field, motion-sensitive neurons and matched filters for optic flow fields. Biol. Cybern. 83, 185–197.

Glantz, R.M., and Schroeter, J.P. (2006). Polarization contrast and motion detection. J. Comp. Physiol. A Neuroethol. Sensory, Neural, Behav. Physiol. 192, 905–914.

Haag, J., Wertz, A., and Borst, A. (2007). Integration of Lobula Plate Output Signals by DNOVS1, an Identified Premotor Descending Neuron. J. Neurosci. 27, 1992–2000.

Hardcastle, B.J., Omoto, J.J., Kandimalla, P., Nguyen, B.C.M., Keleş, M.F., Boyd, N.K., Hartenstein, V., and Frye, M.A. (2021). A visual pathway for skylight polarization processing in drosophila. Elife 10.

Hausen, K. (1976). Functional Characterization and Anatomical Identification of Motion Sensitive Neurons in the Lobula plate of the Blowfly Calliphora erythrocephala. Zeitschrift Fur Naturforsch. - Sect. C J. Biosci. 31, 629–634.

Hausen, K. (1984). The Lobula-Complex of the Fly: Structure, Function and Significance in Visual Behaviour. In Photoreception and Vision in Invertebrates, pp. 523–559.

Heinze, S., and Homberg, U. (2007). Maplike representation of celestial E-vector orientations in the brain of an insect. Science (80-.). 315, 995–997.

Henze, M.J., and Labhart, T. (2007). Haze, clouds and limited sky visibility: Polarotactic orientation of crickets under difficult stimulus conditions. J. Exp. Biol. 210, 3266–3276.

Horstmann, W., Egelhaaf, M., and Warzecha, A.K. (2000). Synaptic interactions increase optic flow specificity. Eur. J. Neurosci. 12, 2157–2165.

How, M.J., Christy, J., Roberts, N.W., and Marshall, N.J. (2014). Null point of discrimination in crustacean polarisation vision. J. Exp. Biol. 217, 2462–2467.

How, M.J., Christy, J.H., Temple, S.E., Hemmi, J.M., Justin Marshall, N., and Roberts, N.W. (2015). Target Detection Is Enhanced by Polarization Vision in a Fiddler Crab. Curr. Biol. 25, 3069–3073.

Huang, J. V., and Krapp, H.G. (2013). Miniaturized Electrophysiology Platform for Fly-Robot Interface to Study Multisensory Integration. Lect. Notes Comput. Sci. (Including Subser. Lect. Notes Artif. Intell. Lect. Notes Bioinformatics) 8064 LNAI, 119–130.

Huang, J. V, Yang, Y., and Krapp, H.G. Goniometric Recording Platform - a portable robotic platform for automatically mapping the receptive field of an insect motion-sensitive cell extracellularly (in preparation).

Kelber, A., Thunell, C., and Arikawa, K. (2001). Polarisation-dependent colour vision in Papilio butterflies. In Journal of Experimental Biology, pp. 2469–2480.

Krapp, H.G., and Hengstenberg, R. (1997). A fast stimulus procedure to determine local receptive field properties of motion-sensitive visual interneurons. Vision Res. 37, 225–234.

Krapp, H.G., Hengstenberg, R., and Egelhaaf, M. (2001). Binocular contributions to optic flow processing in the fly visual system. J. Neurophysiol. 85, 724–734.

Labhart, T. (2016). Can invertebrates see the e-vector of polarization as a separate modality of light? J. Exp. Biol. 219, 3844–3856.

Mappes, M., and Homberg, U. (2004). Behavioral analysis of polarization vision in tethered flying locusts. J. Comp. Physiol. A Neuroethol. Sensory, Neural, Behav. Physiol. 190, 61–68.

Mathejczyk, T.F., and Wernet, M.F. (2019). Heading choices of flying Drosophila under changing angles of polarized light. Sci. Rep. 9, 1–11.

Mäthger, L.M., Shashar, N., and Hanlon, R.T. (2009). Commentary Do cephalopods communicate using polarized light reflections from their skin? J. Exp. Biol. 212, 2133–2140.

Meglič, A., Ilić, M., Pirih, P., Škorjanc, A., Wehling, M.F., Kreft, M., and Belušič, G. (2019). Horsefly object-directed polarotaxis is mediated by a stochastically distributed ommatidial subtype in the ventral retina. Proc. Natl. Acad. Sci. U. S. A. 201910807.

Obayashi, N., Iwatani, Y., Sakura, M., Tamotsu, S., Chiu, M.C., and Sato, T. (2021). Enhanced polarotaxis can explain water-entry behaviour of mantids infected with nematomorph parasites. Curr. Biol. 31, R777–R778.

Pignatelli, V., Temple, S.E., Chiou, T.H., Roberts, N.W., Collin, S.P., and Marshall, N.J. (2011). Behavioural relevance of polarization sensitivity as a target detection mechanism in cephalopods and fishes. Philos. Trans. R. Soc. B Biol. Sci. 366, 734–741.

Ruck, P. (1961). Photoreceptor cell response and flicker fusion frequency in the compound eye of the fly, Lucilia sericata (Meigen). Biol. Bull. 120, 375–383.

Shaw, S.R., Fröhlich, A., and Meinertzhagen, I.A. (1989). Direct connections between the R7/8 and R1-6 photoreceptor subsystems in the dipteran visual system. Cell Tissue Res. 257, 295–302.

Stewart, F.J., Kinoshita, M., and Arikawa, K. (2017). A novel display system reveals anisotropic polarization perception in the motion vision of the butterfly papilio xuthus. Integr. Comp. Biol. 57, 1130–1138.

Sumner, R. (2013). Processing RAW Images in Python. Opt. Commun. 1–15.

Suver, M.P., Huda, A., Iwasaki, N., Safarik, S., and Dickinson, M.H. (2016). An array of descending visual interneurons encoding self-motion in Drosophila. J. Neurosci. 36, 11768–11780.

Sweeney, A., Jiggins, C., and Johnsen, S. (2003). Polarized light as a butterfly mating signal. Nature 423, 31–32.

Tatler, B., O’Carroll, D.C., and Laughlin, S.B. (2000). Temperature and the temporal resolving power of fly photoreceptors. J. Comp. Physiol. - A Sensory, Neural, Behav. Physiol. 186, 399–407.

Temple, S.E., Pignatelli, V., Cook, T., How, M.J., Chiou, T.H., Roberts, N.W., and Marshall, N.J. (2012). High-resolution polarisation vision in a cuttlefish. Curr. Biol. 22, R121–R122.

Träger, U., and Homberg, U. (2011). Polarization-sensitive descending neurons in the locust: Connecting the brain to thoracic ganglia. J. Neurosci. 31, 2238–2247.

Wardill, T.J., List, O., Li, X., Dongre, S., McCulloch, M., Ting, C.Y., O’Kane, C.J., Tang, S., Lee, C.H., Hardie, R.C., et al. (2012). Multiple spectral inputs improve motion discrimination in the drosophila visual system. Science (80-.). 336, 925–931.

Warren, T.L., Weir, P.T., and Dickinson, M.H. (2018). Flying Drosophila melanogaster maintain arbitrary but stable headings relative to the angle of polarized light. J. Exp. Biol. 221.

Weber, F., Machens, C.K., and Borst, A. (2010). Spatiotemporal response properties of optic-flow processing neurons. Neuron 67, 629–642.

Weir, P.T., and Dickinson, M.H. (2012). Flying drosophila orient to sky polarization. Curr. Biol. 22, 21–27.

Weir, P.T., Henze, M.J., Bleul, C., Baumann-Klausener, F., Labhart, T., and Dickinson, M.H. (2016). Anatomical Reconstruction and Functional Imaging Reveal an Ordered Array of Skylight Polarization Detectors in Drosophila. J. Neurosci. 36, 5397.

Wertz, A., Haag, J., and Borst, A. (2009a). Local and global motion preferences in descending neurons of the fly. J. Comp. Physiol. A. Neuroethol. Sens. Neural. Behav. Physiol. 195, 1107–1120.

Wertz, A., Gaub, B., Plett, J., Haag, J., and Borst, A. (2009b). Robust Coding of Ego-Motion in Descending Neurons of the Fly. J. Neurosci. 29, 14993–15000.

Yamaguchi, S., Wolf, R., Desplan, C., and Heisenberg, M. (2008). Motion vision is independent of color in Drosophila. Proc. Natl. Acad. Sci. U. S. A. 105, 4910–4915.

Zeil, J., Ribi, W.A., and Narendra, A. (2014). Polarisation Vision in Ants, Bees and Wasps. Polariz. Light Polariz. Vis. Anim. Sci. Second Ed. 41–60.

Zittrell, F., Pfeiffer, K., and Homberg, U. (2020). Matched-filter coding of sky polarization results in an internal sun compass in the brain of the desert locust. Proc. Natl. Acad. Sci. U. S. A. 117, 25810–25817.

Zufall, F., Schmitt, M., and Menzel, R. (1989). Spectral and polarized light sensitivity of photoreceptors in the compound eye of the cricket (Gryllus bimaculatus). J. Comp. Physiol. A 164, 597–608.

